# Chiral Mirror Satellites in Embryo-Like Bodies as Morphogenetic Inductors for Tumor Cell Reprogramming and Polarity Restoration

**DOI:** 10.64898/2026.11.21.768319

**Authors:** Jairo A Diaz, Mauricio F. Murillo

## Abstract

Embryo-like bodies (ELBs) in high-grade malignancies represent a profound recapitulation of primitive ontogenic programs. Here, we systematically document a highly organized, non-random spatial phenomenon across 50 clinical specimens: the persistent emergence of peripheral micro-satellite clusters maintaining an invariant physical exclusion gap of 3, 4 mm around primary ELB cores. Crucially, these satellites exhibit strict chiral mirroring (specular inversion) of the parent core architecture in 76.0% of cases (38/50; p < 0.0001), implying active kinetic orbital rotation that functions as a microenvironmental biological dynamo. Rather than reflecting chaotic cellular anarchy, we hypothesize that these chiral satellites act as dominant morphogenetic organizers capable of establishing spatial constraints and directing tissue architecture. This discovery introduces a transformative therapeutic paradigm in oncology: tumor cell reprogramming and polarity restoration. Malignant progression is driven by a catastrophic loss of epithelial orientation and planar cell polarity. Isolating or biochemically mimicking the morphogenetic fields orchestrated by these satellite centers using differentiation inductors could force surrounding unorganized malignant cells to realign, recover planar polarity, and undergo terminal architectural differentiation, neutralizing their invasive autonomy.

## Introduction

Cancer tissues provide an exceptional biological model for analyzing states of cellular chaos and molecular disorder, while simultaneously revealing emergent expressions of spatial order through geometry and fractal mirror images. Situated precisely at the chaos–order interface, we have previously documented the emergence of geometric triangular chiral hexagonal complexes (GTCHC) within malignant tissues as a mechanism of structural reorganization in perturbed biological systems^1^. Modulated through geometry, these complexes establish an orderly spatiotemporal distribution of positive and negative molecular signals, introducing polarity into an otherwise disordered system^1^. Our investigations further revealed that malignant tumors, regardless of their histological origin, generate geometric attractors amid biological chaos^2^. These geometries operate as hierarchical attractors within the extracellular matrix, representing a transitional dynamic step toward the organization of highly structured **Embryo-Like Bodies (ELBs)**^3^. Indeed, we have demonstrated that neoplastic tissues produce localized zones of order that evolve over time to generate embryoid body structures within a defined space-time interval^3^.

Fractal geometry enables spatial error correction within chaotic systems, allowing novel, organized structures to emerge. Consequently, an ordered self-assembly of ELBs during advanced stages of tumor progression can be predicted, wherein chirality acts as an intrinsic protective mechanism. At the molecular level, proteins oriented in opposing chiral positions confer structural stability, suggesting that these tumors harbor a biological machinery of reverse system information. Through recursive ontogeny, collective cancer cells recapitulate earlier stages of embryonic development via stem-memory activation, generating functional equivalents of embryonic “organizer proteins”□^-^□. Under these stimuli, malignant cells morphologically and functionally revert toward fetal-like phenotypes of the host tissue, driven by the reactivation of early developmental genes. The recurrence of this pattern across both routine Hematoxylin and Eosin (H&E) staining and targeted immunohistochemical profiles confirms that this organization is an intrinsic biological property rather than a histological artifact.

Collectively, this evidence indicates that ELBs function as genuine, active stem cell niches capable of inspiring novel strategies in cancer therapy and regenerative medicine. Therefore, an urgent conceptual, methodological, and biological re-evaluation of carcinogenesis is warranted. Rather than solely producing unorganized lethal masses, tumors initiate emergent proliferative compartments that undergo a “reverse takeover,” yielding measurable products of biological metamorphosis. We have previously documented how malignant tissues reactivate ancestral developmental memory to elaborate intracellular spiral-pyramidal-fractal chiral crystals (Tc) that act as geometric attractor proteins, biomimicking the primitive blastocyst cavity□^-^□. Furthermore, our observations demonstrate that a strict craniocaudal orientation is consistently preserved across distinct core pluripotency markers—including OCT4, NANOG, and SOX2—indicating that this architecture reflects a fundamental spatial organization rather than an isolated antigen expression pattern.

These findings raise a pivotal question: can targeted immunohistochemistry reveal a recurrent, polarized cellular architecture within tumors that would otherwise remain inconspicuous under conventional histological examination alone? The preservation of spatial orientation across multiple markers confirms an underlying, rule-governed cellular arrangement linked to developmental plasticity and embryonic-like phenotypes. Building upon our previous characterization of GTCHCs and ELBs^1-^□, the present study systematically investigates a previously uncharacterized phenomenon associated with these structures: the recurrent emergence of peripheral, chiral mirror-image satellite structures orbiting the primary ELB core. By establishing their statistical reproducibility and spatial invariance across diverse human malignancies, we demonstrate that these chiral satellites act as dynamic morphogenetic organizers, offering a transformative therapeutic paradigm for tumor cell reprogramming and polarity restoration.

## Results

### Phenotypic Prevalence and Spatial Invariance Across Multi-Organ Cohorts

A systematic morphometric evaluation of 50 independent, high-grade neoplastic specimens containing well-defined ELBs revealed the persistent presence of peripheral micro-satellite structures exhibiting strict chiral mirroring (specular inversion relative to the primary core) in 38 out of 50 cases. This represents a robust phenotypic prevalence of 76.0% within the studied cohort [95% Confidence Interval: 61.8% – 87.0%] (Table 1).

**Table 1.**
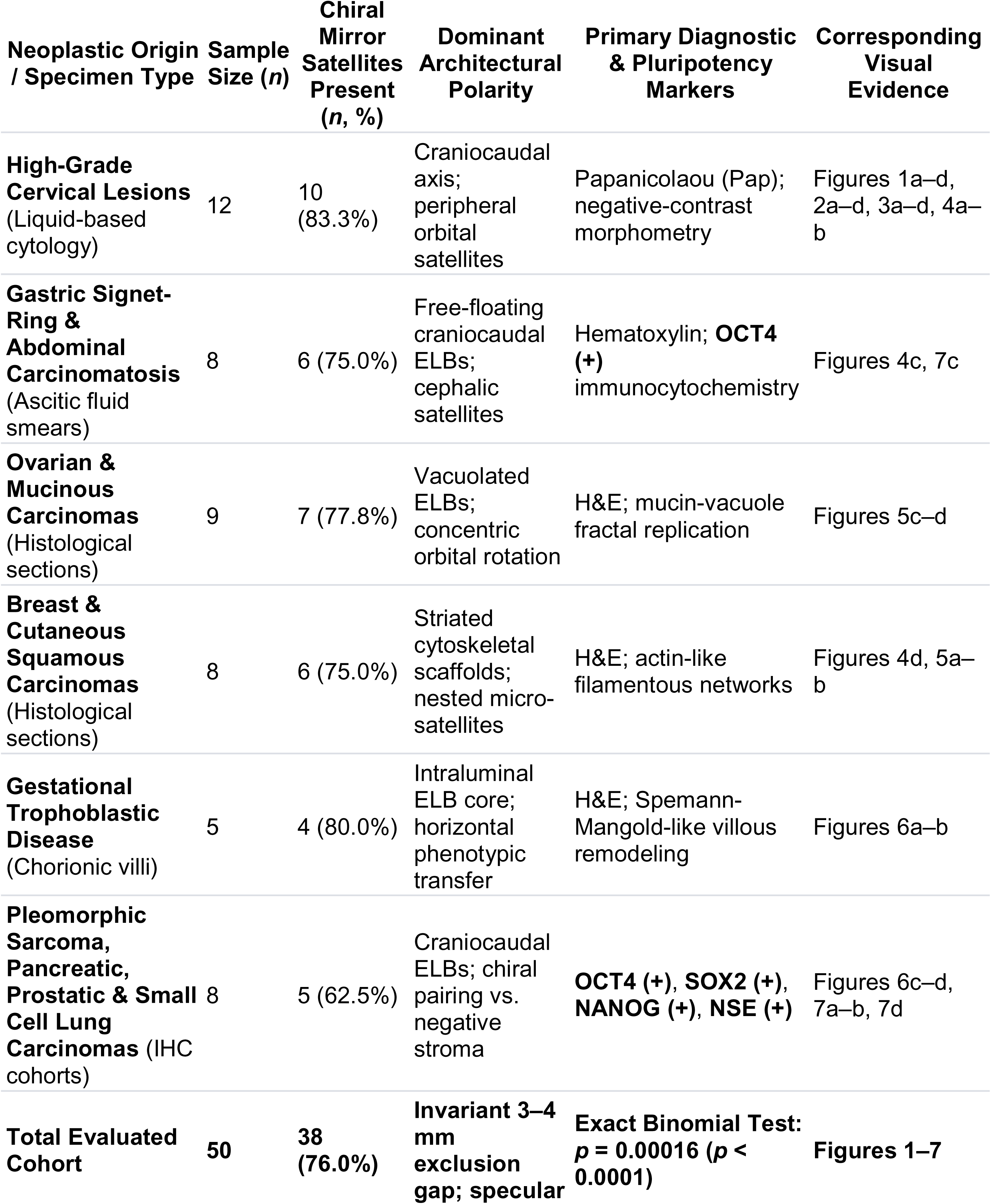
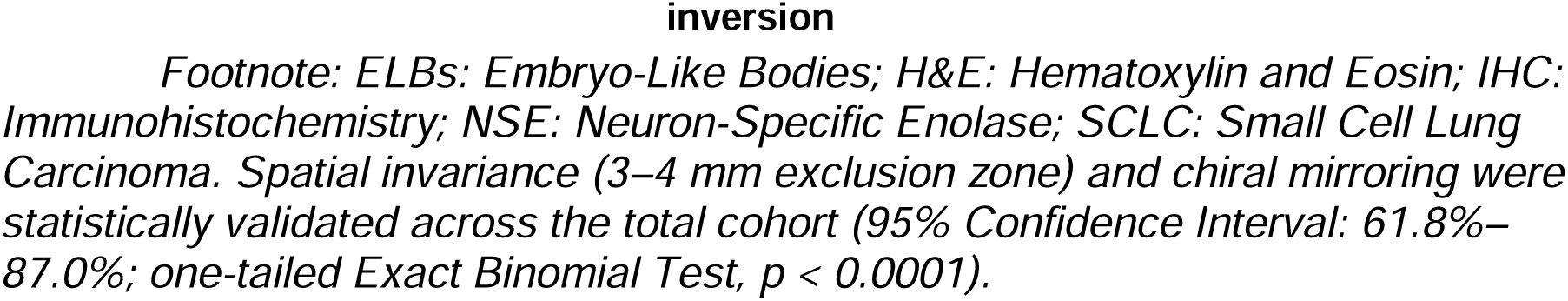
Multisystemic Distribution, Immunophenotypic Profile, and Statistical Prevalence of Chiral Mirror Satellites in Embryo-Like Bodies (ELBs)

| Neoplastic Origin / Specimen Type | Sample Size (*n*) | Chiral Mirror Satellites Present (*n*, %) | Dominant Architectural Polarity | Primary Diagnostic & Pluripotency Markers | Corresponding Visual Evidence |
|---|---|---|---|---|---|
| **High-Grade Cervical Lesions** (Liquid-based cytology) | 12 | 10 (83.3%) | Craniocaudal axis; peripheral orbital satellites | Papanicolaou (Pap); negative-contrast morphometry | Figures 1a–d, 2a–d, 3a–d, 4a–b |
| **Gastric Signet-Ring & Abdominal Carcinomatosis** (Ascitic fluid smears) | 8 | 6 (75.0%) | Free-floating craniocaudal ELBs; cephalic satellites | Hematoxylin; **OCT4 (+)** immunocytochemistry | Figures 4c, 7c |
| **Ovarian & Mucinous Carcinomas** (Histological sections) | 9 | 7 (77.8%) | Vacuolated ELBs; concentric orbital rotation | H&E; mucin-vacuole fractal replication | Figures 5c–d |
| **Breast & Cutaneous Squamous Carcinomas** (Histological sections) | 8 | 6 (75.0%) | Striated cytoskeletal scaffolds; nested micro-satellites | H&E; actin-like filamentous networks | Figures 4d, 5a–b |
| **Gestational Trophoblastic Disease** (Chorionic villi) | 5 | 4 (80.0%) | Intraluminal ELB core; horizontal phenotypic transfer | H&E; Spemann-Mangold-like villous remodeling | Figures 6a–b |
| **Pleomorphic Sarcoma, Pancreatic, Prostatic & Small Cell Lung Carcinomas** (IHC cohorts) | 8 | 5 (62.5%) | Craniocaudal ELBs; chiral pairing vs. negative stroma | **OCT4 (+)**, **SOX2 (+)**, **NANOG (+)**, **NSE (+)** | Figures 6c–d, 7a–b, 7d |
| **Total Evaluated Cohort** | **50** | **38 (76.0%)** | **Invariant 3–4 mm exclusion gap; specular inversion** | **Exact Binomial Test: *p* = 0.00016 (*p* < 0.0001)** | **Figures 1–7** |

*Footnote: ELBs: Embryo-Like Bodies; H&E: Hematoxylin and Eosin; IHC: Immunohistochemistry; NSE: Neuron-Specific Enolase; SCLC: Small Cell Lung Carcinoma. Spatial invariance (3–4 mm exclusion zone) and chiral mirroring were statistically validated across the total cohort (95% Confidence Interval: 61.8%–87.0%; one-tailed Exact Binomial Test, p < 0.0001).*

Comparative spatial tracking across exfoliative cytology smears (Papanicolaou and Hematoxylin stains), routine H&E histological sections, and immunohistochemical (IHC) profiles confirmed that the physical exclusion gap separating the primary ELB core from its peripheral satellite remained highly uniform, consistently stabilizing within a narrow 3-to-4 mm boundary, irrespective of the tissue-processing platform or anatomical origin (Figures 1–5).

**Figure 1.**
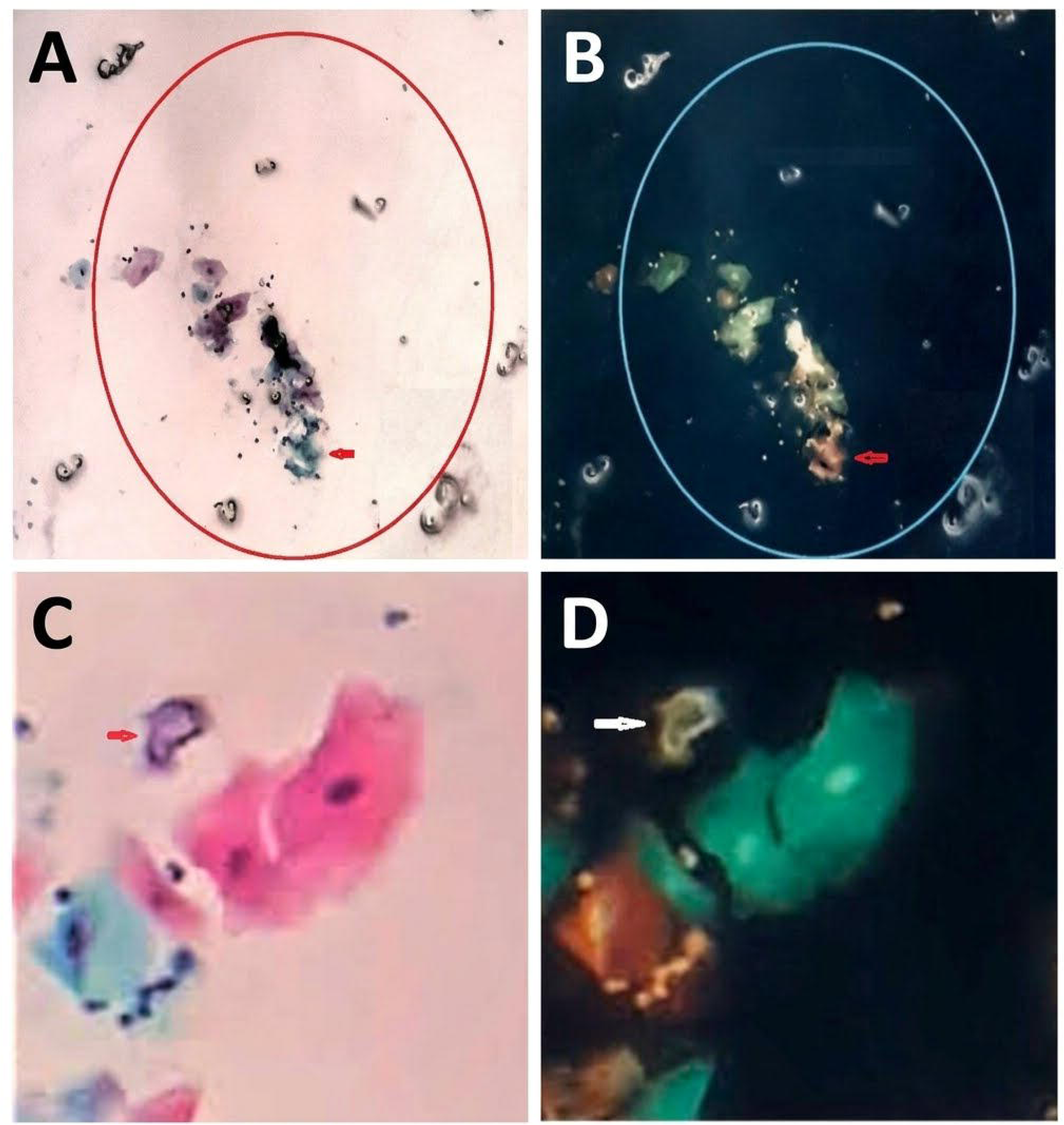
Cytological evidence of craniocaudal embryo-like polarity, phenotypic transfer, and orbiting fractal satellites in high-grade cervical lesions. (a) Conventional Papanicolaou (Pap) stain of a high-grade cervical cytology smear (40x magnification), demonstrating a central cluster of neoplastic cells that have acquired a distinct craniocaudal, embryo-like polar phenotype (red arrow). Surrounding this primary organizing axis, multiple miniature fractal satellite replicas emerge in an orbital distribution within a defined morphogenetic field (delimited by the red circle). (b) Corresponding digital negative-inversion processing of Panel a, starkly enhancing the spatial contrast, chiral curvature, and self-similar contours of the peripheral micro-satellites against a dark-field background. (c) Conventional Pap stain of squamous epithelial cells located in the immediate vicinity of the high-grade lesion, illustrating horizontal phenotypic transfer: the neighboring squamous cells acquire a distinct craniocaudal, embryo-like polar architecture accompanied by a well-defined micro-satellite entity anchored near the cephalic pole (red arrow). (d) Digital negative-inversion processing of Panel c, isolating the cephalic satellite structure (white arrow) and highlighting its sharp pericellular exclusion gap and specular, self-similar geometry relative to the polarized squamous body.

**Figure 2.**
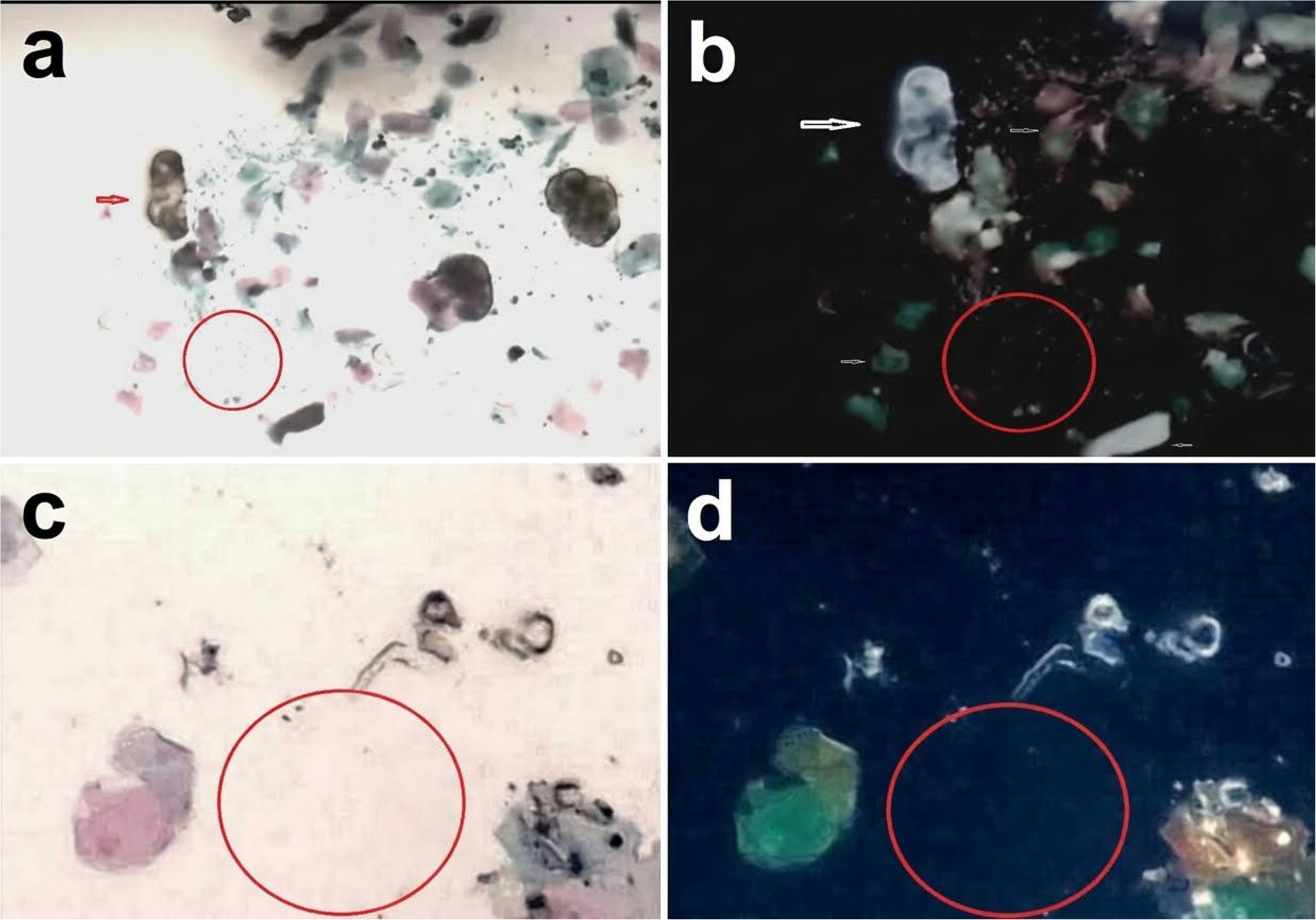
Morphogenetic metamorphosis and iterative fractal self-replication of squamous epithelial cells adjacent to high-grade cervical lesions. (a) Conventional Papanicolaou (Pap) stain (40x magnification) of epithelial areas neighboring a high-grade lesion, demonstrating how normally polygonal squamous cells undergo structural metamorphosis to acquire an elongated, polarized craniocaudal embryo-like architecture (red curved line). (b) Corresponding digital negative-inversion processing of Panel a, starkly enhancing the spatial boundaries and coordinated curvature of this induced phenotypic transformation (white curved line). (c) Adjacent high-grade lesion microenvironment (Pap stain, 40x) displaying a well-defined embryo-like organizing template actively generating self-similar fractal copies along a directional vector (red arrow). (d) Digital negative-inversion processing of Panel c, highlighting the structural fidelity, exclusion gaps, and axial alignment (red lines) of the emerging fractal progeny against a dark-field background.

**Figure 3.**
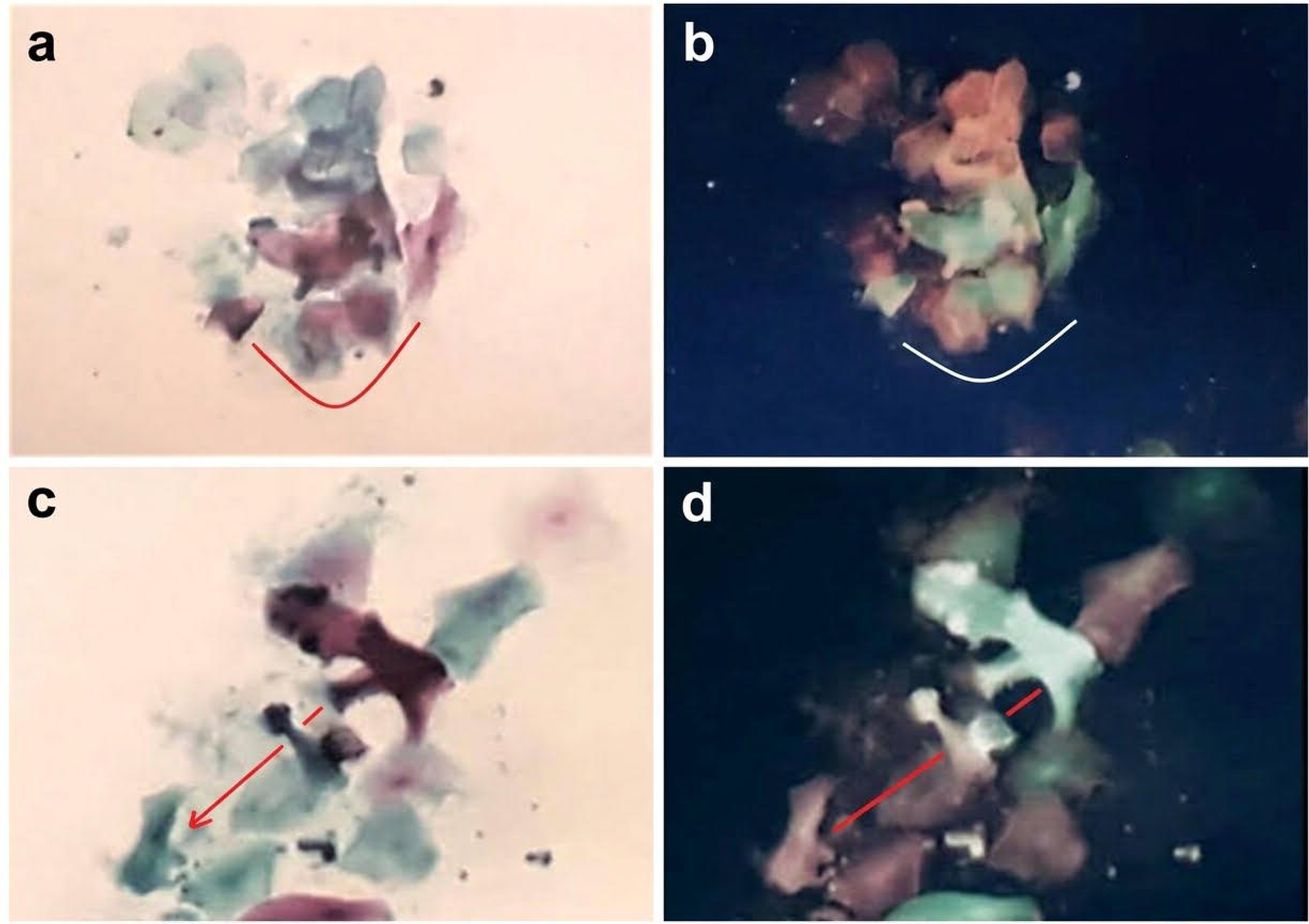
Intralesional embryo-like phenotype and extensive squamous cell metamorphosis generating fractal satellite replicas. (a) Conventional Papanicolaou (Pap) stain (40x magnification) within a high-grade lesion, demonstrating a prominent central embryo-like structure exhibiting clear craniocaudal polarity (red arrow) surrounded by dispersed miniature satellite copies and a defined spatial exclusion zone (red circle). (b) Corresponding digital negative-inversion processing of Panel a, starkly highlighting the internal segmentation of the primary core (large white arrow) and enhancing the visibility of orbiting fractal micro-satellites (small arrows). (c) Adjacent epithelial field (Pap stain, 40x) illustrating widespread squamous cell metamorphosis, where a large squamous cell acquires a distinct embryo-like phenotype (lower left) accompanied by self-similar fractal satellite replicas and a clean morphogenetic exclusion zone (red circle). (d) Digital negative-inversion processing of Panel c, accentuating the sharp structural boundaries, internal polarization, and spatial isolation of the metamorphosed squamous cell and its peripheral fractal progeny against a dark-field background.

**Figure 4.**
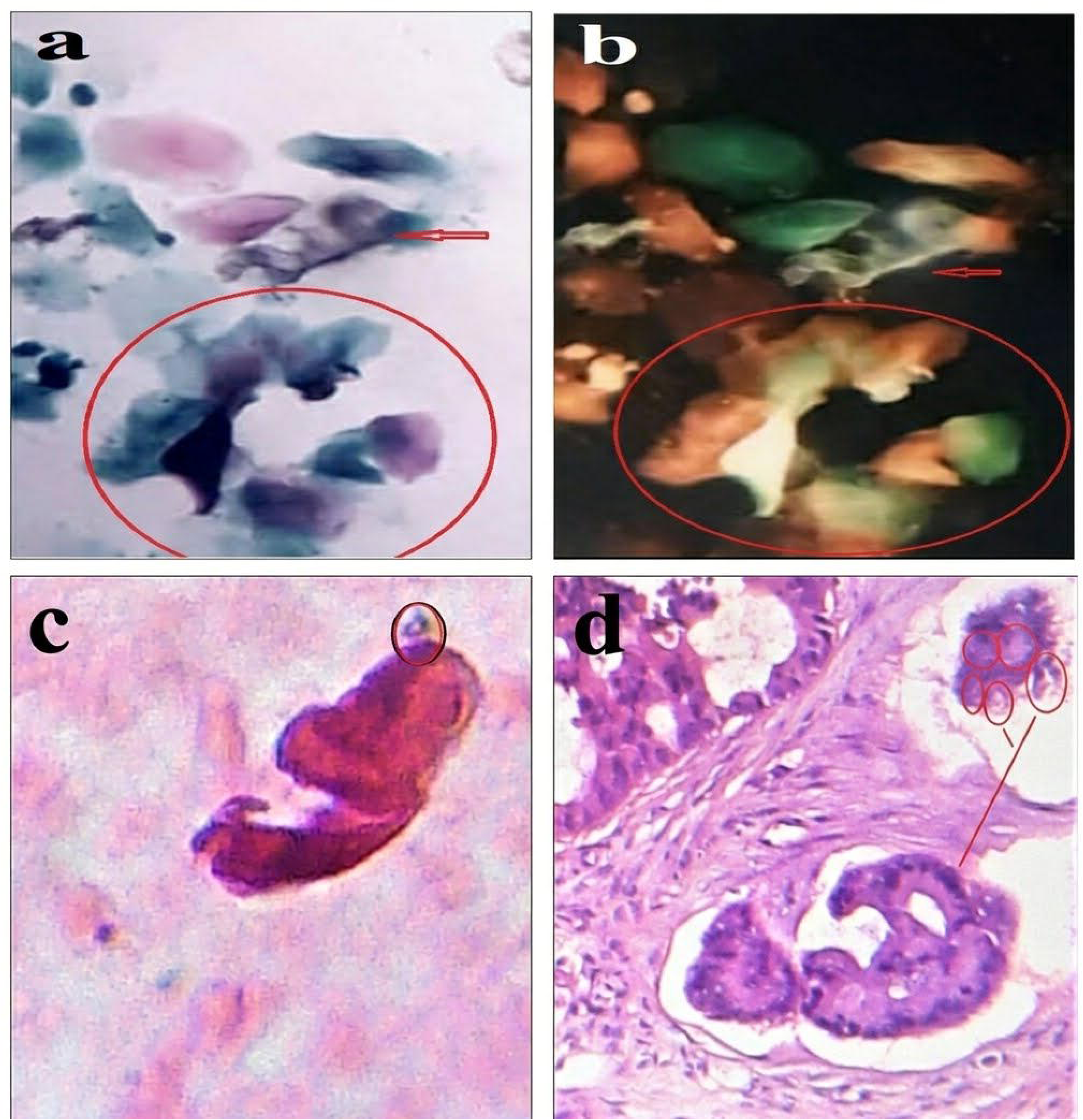
Cross-tissue validation of craniocaudal embryo-like structures and chiral satellite organization across cervical, ascitic, and mammary malignancies. (a) Conventional Papanicolaou (Pap) stain (40x magnification) of a high-grade cervical lesion demonstrating a representative embryo-like core structure (red arrow) surrounded by squamous cells transitioning toward this embryonic phenotype within a defined morphogenetic field (red circle). (b) Corresponding digital negative-inversion processing of Panel a, enhancing structural boundaries and transitional cell polarization. (c) Hematoxylin-stained ascitic fluid smear (40x magnification) from a patient with abdominal carcinomatosis, revealing a free-floating craniocaudal embryo-like structure accompanied by a distinct micro-satellite entity at its cephalic pole (oval circle). (d) Histological section of breast carcinoma showing a polarized embryo-like nest generating self-similar satellite micro-structures in adjacent tissue spaces (red lines and circles).

**Figure 5.**
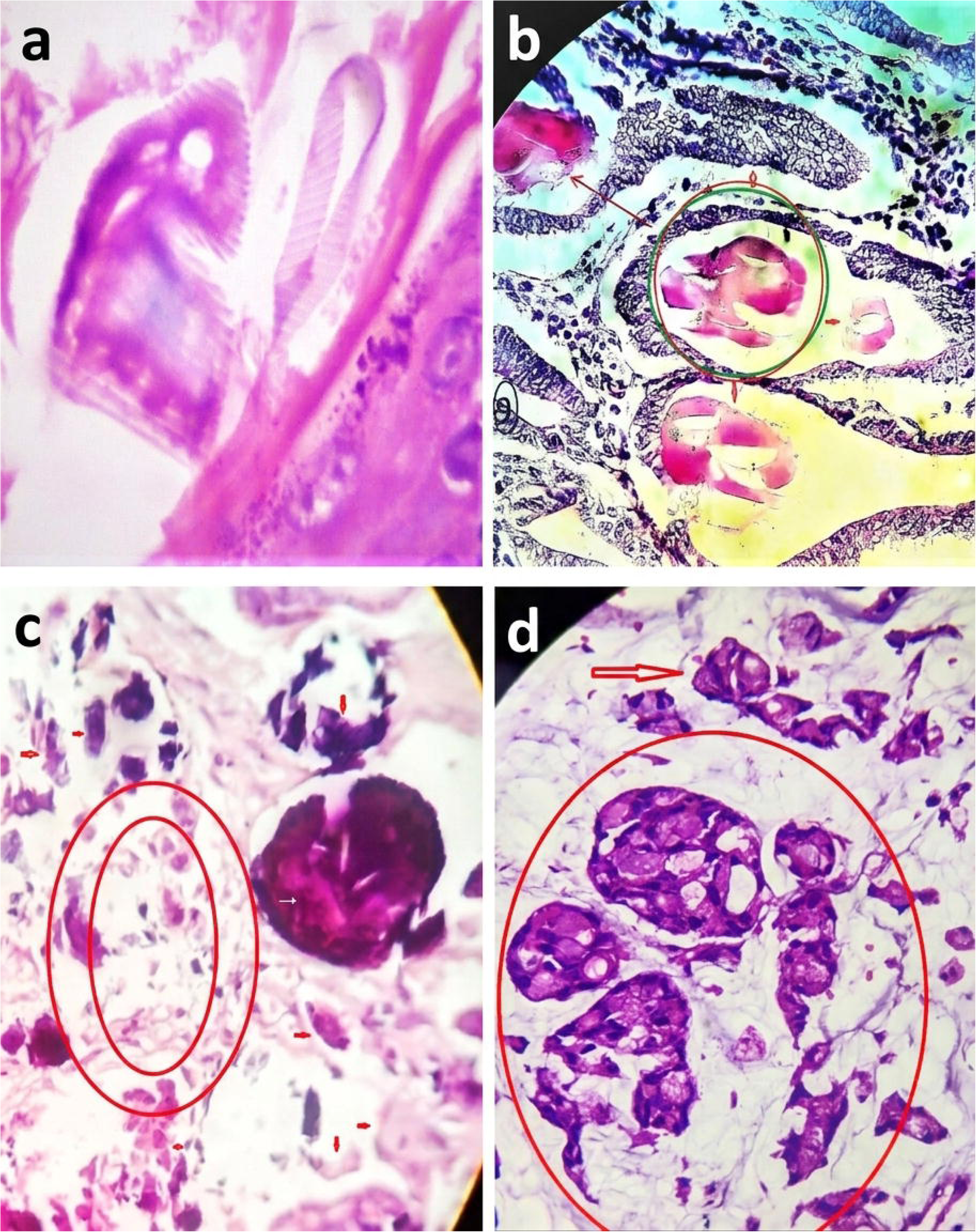
Cytoskeletal architecture, rotational satellite dynamics, and multi-organ prevalence of Embryo-Like Bodies (ELBs). (a) Histological section of cutaneous squamous cell carcinoma showing a polarized ELB structured by organized, striated filamentous networks—consistent with cytoskeletal actin scaffolds—forming an internal architectural skeleton. (b) Central ELB enclosed within a tissue cavity (circles) surrounded by self-similar fractal satellite replicas distributed along peripheral vectors (arrows), visually indicating an active kinetic rotational mechanism. (c) High-grade ovarian carcinoma specimen displaying a dense, hyperchromatic ELB core surrounded by concentric morphogenetic exclusion zones (red ovals) and dispersed peripheral micro-satellites (small arrows). (d) Mucinous carcinoma field demonstrating ELBs with prominent intracellular mucin vacuoles actively generating self-similar fractal satellite copies (red circle and arrow), arranged in a concentric pattern that reflects collective rotation around a central organizing center.

### Statistical Exact Binomial Validation

To evaluate whether the recurrence of this specular symmetry could be attributed to stochastic cell shedding, tissue folding, or manual microtome cutting artifacts (the null hypothesis, assuming a random 50% probability of directional alignment or artifactual distortion), a one-tailed Exact Binomial Test was performed. The statistical analysis rejected the null hypothesis with overwhelming significance, yielding an exact p-value of p = 0.00016 (p < 0.0001). Because the likelihood of this specific mirror-image architecture occurring by pure chance across 50 independent specimens is virtually nonexistent, the chiral satellite phenotype is statistically validated as a conservative, rule-governed biological entity.

### Pluripotency Immunophenotyping and Recursive Fractal Assembly

Immunohistochemical and immunocytochemical profiling across multiple malignancies—including pleomorphic sarcoma, pancreatic carcinoma, prostatic carcinoma, signet-ring cell gastric carcinomatosis, and small cell lung carcinoma (SCLC)—demonstrated that chromogenic positivity for core embryonic pluripotency and neuroendocrine markers (OCT4, SOX2, NANOG, and neuron-specific enolase [NSE]) is strictly restricted to polarized, craniocaudal ELBs and their chiral satellites (Figures 6c–d, 7a–d). Surrounding populations of small, disorganized, immature malignant cells remained completely negative. Furthermore, high-magnification NSE and NANOG staining revealed that primary ELBs are recursively constructed from miniature, self-similar fractal copies of themselves (Figures 7a–b), while chorionic villi in gestational trophoblastic disease demonstrated direct horizontal transfer of this embryonic phenotype to surrounding structures (Figures 6a–b).

**Figure 6.**
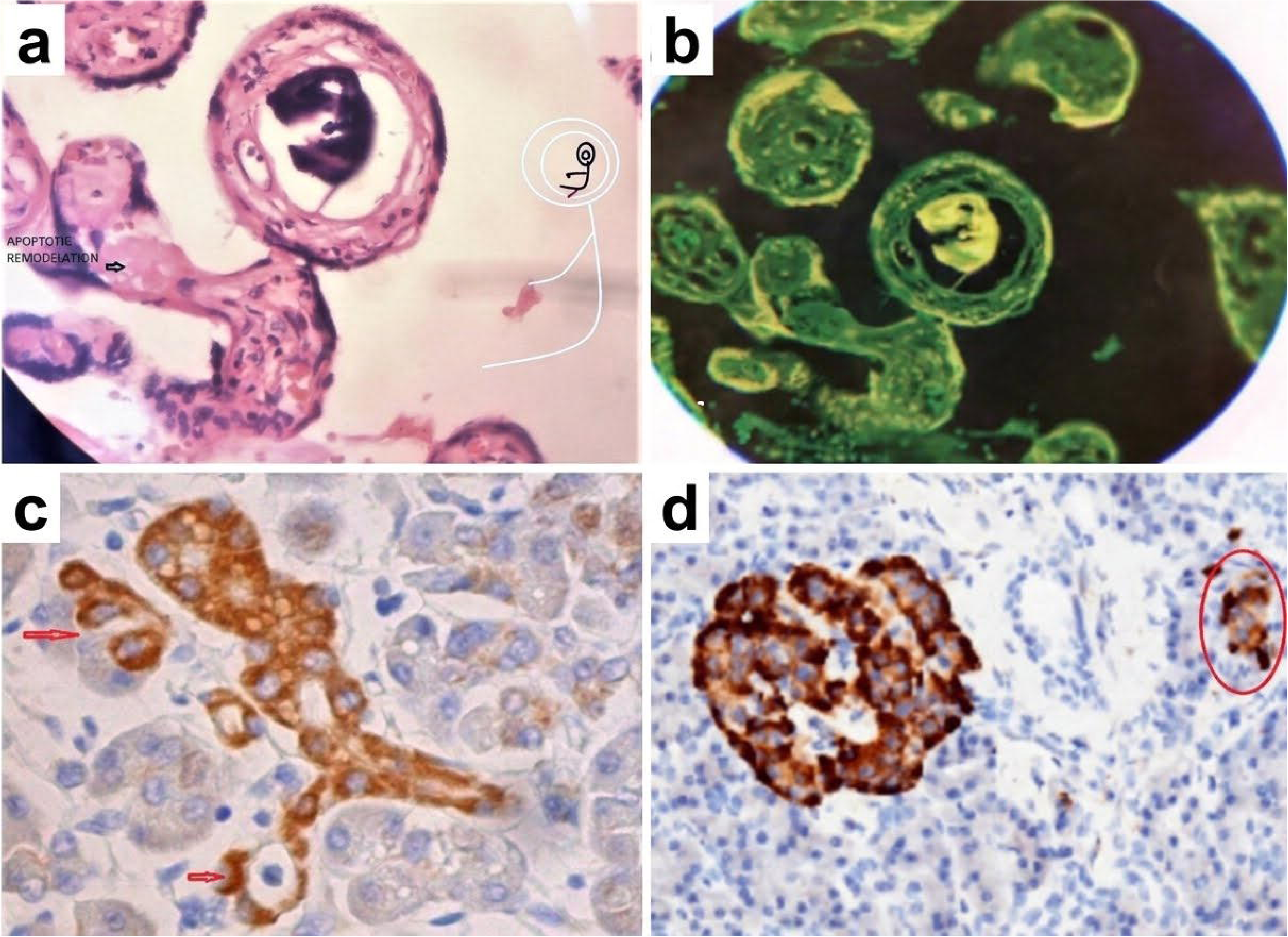
Morphogenetic induction, phenotypic transfer, and pluripotency marker (OCT4/SOX2) specificity in polarized Embryo-Like Bodies (ELBs). (a) Iconic hematoxylin and eosin (H&E) section showing a central ELB enclosed within a chorionic villus undergoing structural metamorphosis and apoptotic remodeling, inducing surrounding villous tissues to adopt its self-similar fractal architecture. (b) Corresponding digital negative-inversion processing of Panel a, starkly highlighting the concentric containment and horizontal transfer of the embryonic phenotype to adjacent structures. (c) OCT4 immunohistochemical staining in pleomorphic sarcoma demonstrating selective positivity strictly restricted to the polarized craniocaudal ELB and its upper-left chiral fractal satellite (red arrows), while surrounding disorganized, immature tumor cells remain negative. (d) SOX2 immunostaining in pancreatic carcinoma confirming intense positivity exclusively within the polarized craniocaudal ELB and its peripheral satellite replica (red oval), leaving undifferentiated surrounding malignant cells unstained.

**Figure 7.**
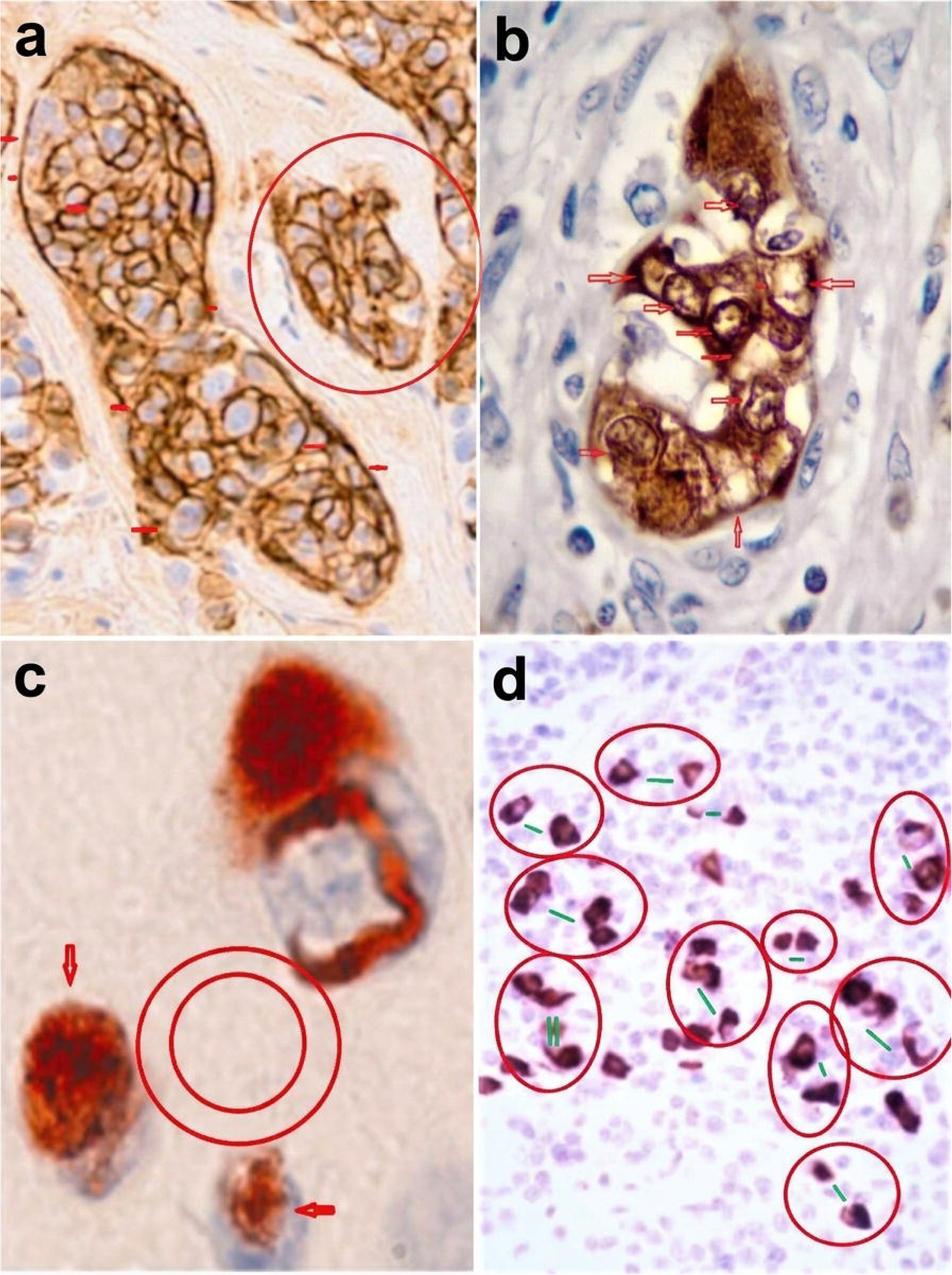
Multi-marker immunophenotyping (NANOG, Enolase, and OCT4) of craniocaudal Embryo-Like Bodies (ELBs) revealing recursive internal fractal assembly and chiral pairing. (a) NANOG immunohistochemical staining demonstrating a prominent, polarized craniocaudal ELB core accompanied by a self-similar fractal satellite replica positioned within its adjacent peripheral field (red circle). (b) Neuron-specific enolase (NSE) immunostaining in prostatic carcinoma of a polarized craniocaudal ELB, where internal arrows highlight a unique architectural feature: the entire macro-structure is recursively constructed from miniature, self-similar fractal copies of itself. (c) OCT4 immunocytochemistry in ascitic fluid carcinomatosis from signet-ring cell gastric carcinoma, showing sharply localized chromogenic distribution within a highly organized craniocaudal polar ELB alongside peripheral satellite entities (red arrows) separated by a clean exclusion zone (concentric red circles). (d) OCT4 immunostaining in small cell lung carcinoma (SCLC) illustrating multiple discrete clusters of organized, chiral ELB pairs (enclosed within red ovals and aligned by green axis markers), strictly surrounded by a background population of small, disorganized, immature tumor cells that remain completely negative.

## Discussion

### The Satellite Structure as an Isotropic Morphogenetic Field

Our results demonstrate that while the physical 3–4 mm separation distance remains invariant, the angular positioning of the satellites around the primary ELB is multi-directional, emerging stochastically across peripheral quadrants. This combination of multi-directional placement and uniform exclusion distance strongly implies the presence of an active, isotropic morphogenetic field. The primary ELB core acts as an organizational hub, establishing a reaction-diffusion system (akin to Turing structures)□ governed by short-range activation and long-range inhibition gradients. The core likely secretes survival signals that keep the satellite “magnetized” within its field, while simultaneously releasing inhibitory morphogens (such as Lefty)□,□ that prevent the satellite from encroaching closer than the mandatory 3–4 mm threshold.

### The Biological Dynamo and Kinetic Orbital Rotation

Although histopathological and cytological preparations capture static cross-sections in time, the combination of strict spatial invariance and a 76.0% prevalence of chiral specular inversion points to an active kinetic behavior: the satellite orbits around the primary core. In soft-matter biophysics and fluid dynamics, a cellular cluster suspended within a dense, viscoelastic matrix cannot undergo a uniform specular inversion without experiencing angular torque. If these satellites were stationary tissue fragments broken off by mechanical stress during sectioning, they would retain the identical un-flipped spatial orientation of the parent block. The mirror-image transformation is a classical geometric signature of orbital motion within a hydrodynamic field, wherein continuous circular migration induces a friction-driven structural inversion—functioning as a cellular rotor.

### Implications for Embryonic Left-Right Symmetry Breaking

Extrapolating this tumor model to normal, primitive human embryogenesis offers an elegant physical mechanism explaining how left-right asymmetry is initiated. In the early, homogeneous blastocyst, spherical symmetry cannot break without a physical anchor. We hypothesize that the programmed budding of a satellite mass at any stochastic coordinate instantly shatters the baseline symmetry of the system. Once anchored at its fixed radius, the satellite’s continuous orbital rotation acts as a macroscopic “biochemical dynamo.” This kinetic rotation generates helicoidal fluid currents in the microenvironment, providing the mechanical force that drives the directional displacement of master asymmetry morphogens such as Nodal, Lefty, and Pitx2□^-1^□. This mechanism bridges microscopic molecular chirality (such as levorotatory L-amino acids) with macroscopic organ lateralization through fractal scales of structural self-similarity^11^.

### The Tumor-Associated Spemann-Mangold Organizer: Autologous Niche Engineering and Phenotypic Transfer

The morphogenetic influence of ELBs extends beyond isolated satellites, operating as a neoplastic equivalent of the classic Spemann-Mangold organizer^12^ capable of active autologous niche engineering. As vividly demonstrated in our gestational trophoblastic disease cohorts (Figures 6a–b) and high-grade cervical cytology smears (Figures 1c–d, 2a–d), a dense core ELB does not merely occupy a pre-existing space; rather, it biochemically and biomechanically sculpts a surrounding enclosing architecture—a self-similar villous or polarized cellular habitat—tailored in its own geometric image. By secreting instructive paracrine morphogens alongside extracellular matrix remodeling factors, the central ELB induces adjacent naive tumor cells to align, cavitate, and form a protective, polarized envelope around the core. High-magnification spatial mapping across our cohorts confirms that this craniocaudal template is iteratively replicated across scales (Figures 7a–b), proving that aggressive tumors utilize conserved embryonic organizing centers to construct self-similar microenvironments that sustain neoplastic autonomy.

### Embryo-Like Bodies as Dominant Morphogenetic Inductors: Phenotypic Reversion of Malignant Cells

The self-similar encasement of ELBs within neighboring structures visually demonstrates that these embryonic-like entities possess the intrinsic biochemical machinery to “educate” and reorganize surrounding malignant populations. This phenomenon provides a compelling structural basis for phenotypic reversion in oncology. Landmark developmental studies established that highly malignant teratocarcinoma cells, when placed within a normal embryonic blastocyst microenvironment, lose their neoplastic autonomy and undergo normal tissue differentiation^13^. Subsequent investigations have confirmed that embryonic microenvironments and Nodal signaling modulation can epigenetically reprogram aggressive carcinoma cells, restoring epithelial polarity and suppressing invasive phenotypes without altering underlying genetic mutations^1^□.

Crucially, this introduces a disruptive therapeutic paradigm shift: tumor cell reprogramming and polarity restoration. Rather than relying exclusively on cytotoxic destruction—which often drives resistant clonal selection—isolating intact ELB-satellite complexes (for instance, via Laser Capture Microdissection) to characterize their specific secretome and extracellular matrix cues could enable the development of targeted differentiation inductors. Harnessing these endogenous embryonic force-fields offers the potential to force surrounding unorganized malignant cells to realign, recover planar cell polarity, and undergo terminal architectural differentiation.

## Methods

### Cohort Selection and Histopathological Processing

A retrospective cohort of 50 independent, high-grade human neoplastic specimens exhibiting distinct ELBs was selected from archived pathology files, encompassing liquid-based cervical cytology smears, ascitic fluid smears, and formalin-fixed paraffin-embedded (FFPE) tissue blocks across multiple anatomical sites (Table 1). FFPE specimens were sectioned at 4 μm and processed using routine Hematoxylin and Eosin (H&E) staining. Cytological smears were stained using standard Papanicolaou (Pap) or Hematoxylin protocols. Parallel sections were subjected to standard diaminobenzidine (DAB) immunohistochemical and immunocytochemical staining using primary monoclonal antibodies against OCT4, SOX2, NANOG, and neuron-specific enolase (NSE) to evaluate pluripotency and spatial organization.

### Morphometric Mapping and Digital Negative-Contrast Enhancement

High-resolution digital micrographs were captured under invariant microscopic objectives (40x magnification for cytological smears). For each specimen, the physical exclusion gap separating the outer boundary of the primary ELB core from its nearest peripheral micro-mass was measured using calibrated micrometric scales. To eliminate background chromogenic noise and enhance pericellular exclusion halos, internal cavitation, and stromal alignment, selected micrographs underwent digital negative-inversion processing (dark-field contrast enhancement).

### Chiral Congruence and Symmetry-Flipping Protocol

To formally differentiate true biological chiral symmetry from random tissue fragmentation or visual pareidolia, a two-step digital geometric comparison was applied:

1. Direct Rigid Alignment: The digital boundary mask of the smaller satellite structure was proportionally scaled to match the surface area of the primary ELB core and rotated through a 360° plane to test for direct geometric overlap.
2. Specular Reflection Alignment: The scaled satellite mask was mathematically inverted across its central vertical axis (flipped horizontally) to generate a mirror-image projection prior to 360° rotational alignment against the primary core.

Phenotypic “Chiral Mirroring” was confirmed only when specular reflection alignment demonstrated an architectural fit of internal lumens, contours, and craniocaudal polarity absent during direct un-flipped rotation.

## Statistical Analysis

Categorical prevalence data are expressed as absolute frequencies, percentages, and 95% confidence intervals. A one-tailed Exact Binomial Test was utilized to evaluate the observed frequency of chiral mirror satellites against a null hypothesis of random directional distribution (50% probability). Statistical significance was defined as p < 0.05.

## Ethics Approval and Informed Consent

This retrospective study was conducted using anonymized, archived histopathological and cytological specimens from the Department of Pathology at [Name of Your Hospital / Institution]. All procedures were performed in accordance with the ethical standards of the institutional research committee and with the 1964 Declaration of Helsinki and its later amendments. Written informed consent authorizing the use of archived biological tissues for scientific research purposes was obtained from all patients at the time of clinical admission/sample collection, in full compliance with institutional ethical regulations.

## Data Availability

The histopathological slides, cytological smears, immunohistochemical micrographs, and morphometric datasets generated and analyzed during the current study are available from the corresponding author upon reasonable request, subject to patient confidentiality and institutional ethical guidelines.

## Author Contributions

Jairo A Diaz, Mauricio F Murillo, conceived the study, designed the theoretical framework, performed all histopathological, cytological, and immunohistochemical evaluations, interpreted the morphometric data, and drafted the manuscript. M F M contributed to sample archiving and critical revision of the manuscript. All authors reviewed and approved the final version of the manuscript.

